# Stretched Exponential Modeling Reveals Drug-Specific Kinetics in Human Engineered Skeletal Muscle

**DOI:** 10.64898/2026.06.08.730797

**Authors:** Mattias Luber, Bruno Schmelz, Christof Lenz, Timo Betz

## Abstract

Understanding muscle tissue is key not only for advancing our knowledge in biophysical function of the human body, but can also open new paths to better description of pathological states, improve treatment and training and even provide insights for drug development. While using excised muscles is an excellent way to study muscular function, this limits access to non-human tissue, as human muscle tissue donation is very limited. The past decade has seen a rise in 3D *in vitro* models of human skeletal muscle, however many current experimental approaches to create such models often struggle with the high variability of primary cells or the limited translational relevance in cases where murine lines are used. Furthermore, the state-of-the-art functional analysis is typically focusing at peak force, which overlooks critical kinetic information. In this study, we present an systematic approach to generate functional engineered skeletal muscle tissues (ESMs), measure the contraction force dynamics and model these with a new approach to extract the effect of a series of standard pharmacological modulators.

Within 14 days, LHCN-M2 ESMs mature into aligned, multinucleated myofibers expressing key sarcomeric markers and exhibiting robust excitation-contraction coupling. To decode these dynamics, we introduce a novel mathematical model based on stretched exponential functions. This framework accurately captures the heterogeneous contraction and relaxation phases across diverse phenotypes using only six interpretable parameters.

We validated the platform’s sensitivity using a library of pharmacological modulators. Our kinetic modeling revealed a distinct parameter fingerprint for different drug classes such as the specific changes of contraction kinetics that peak force alone could not detect. Additionally the presence of the necessary drug targets as well as the state of maturation were investigated by proteomic profiling. Together, study provides a scalable, high-fidelity human platform for high-content pharmacological screening and muscle biophysics.

## INTRODUCTION

Engineered skeletal muscle tissues (ESMs) have emerged as a paradigm-shifting tool in translational biomedical research, offering in vitro models that closely recapitulate the native tissue microenvironment, architecture, and mechanical behavior. Unlike traditional two-dimensional (2D) monolayer cultures, 3D ESMs facilitate a higher degree of maturation through prolonged culture periods and provide increased physiological relevance via interactions with an extracellular matrix (ECM) (1–3). Crucially, these systems enable the functional measurement of contractile force through pillar-based platforms, offering a direct window into the mechanical state of the tissue.

In the context of translational research, the choice of tissue generating cells is a critical determinant of both physiological relevance and reproducibility. Historically, 3D muscle models have relied on immortalized murine cell lines, such as C2C12 (4–7) or L6 (8). While these lines are robust and well-characterized, their utility in understanding human muscle is often limited by significant inter-species differences that hinder the translation of results to human clinical outcomes. To bridge this gap, recent efforts have shifted toward human primary myoblasts(1, 9–13) or induced pluripotent stem cells (iPSCs) (14–16). Although primary cells offer high clinical fidelity (9), they suffer from low proliferation rates and significant donor-to-donor variability, which compromises reproducibility. Conversely, while iPSC-based protocols can yield high consistency, they are often complex, lengthy, and technically demanding (17–19). Human immortalized cell lines could represent a potential “sweet spot,” providing human-derived genetics with the high consistency and scalability of a cell line. However, to date, only a few human immortalized lines, such as AB1167, have been investigated in 3D (20–22), and their limited availability remains a significant barrier to widespread adoption.

Beyond tissue engineering, the quantitative interpretation of contractile properties remains a significant challenge in functional profiling. The modeling of muscle dynamics has a storied history, ranging from classic Hill-type (23) to more complex Huxley-type (24) cross-bridge models. While multi-scale models provide extensive biophysical insights, their implementation frequently requires complex systems of differential equations and numerous parameters that are experimentally underdetermined. To get the relevant working parameters specialized measurements, such as molecular binding rates (25, 26) are necessary. As a result, many application-oriented studies default to “Peak Force” as the primary readout for functional performance (6, 27, 28). However, peak force is a one-dimensional metric that fails to capture the full temporal dynamics of the contraction-relaxation cycle-dynamics that often encode dynamical features that are relevant to describe the force generation. This is in particular important when studying a specific mechanism of pharmacological modulators or the underlying integrity of the cytoskeletal architecture.

In this study, we present an integrated approach designed to address these challenges of scalability, reproducibility, and analytical depth. Our main contributions are threefold:

- We demonstrate the suitability of the widely available human immortalized myoblast cell line, LHCN-M2 (29), for generating functional 3D skeletal muscle tissues. We show that these tissues express key differentiation markers, exhibit spontaneous contractions, and demonstrate robust electrical excitability.
- We demonstrate the potential of our previously introduced platform (6) to study biophysical parameters of muscle tissue using a library of pharmacological modulators, including myosin inhibitors, troponin activators, and cytoskeletal disruptors.
- We propose a compact yet powerful mathematical model based on stretched exponential functions. We demonstrate that this model accurately captures the kinetics of tetanic contractions across a broad spectrum of phenotypes using a minimal set of interpretable parameters, providing a more granular high-content readout than peak force alone.

## RESULTS

### LHCN-M2 cells differentiate into mature and functional 3D muscle tissues

To investigate the contractile dynamics of ESMs we generated 3D microtissues using the immortalized human myogenic cell line LHCN-M2(29). The tissues were raised and cultured within a previously published pillar-base plattform (MicroPlate) (6), that consist of an 8-Well injection-molded polystyrene base and a CNC-milled PMMA-lid featuring two cantilever-pillars per well (Figure 1). To prevent the attachment of the tissue to the surface material of the wells, the Microplate was coated with a 5% Pluronic solution prior to seeding the tissues, to produce a surface that is non-adhesive to cells.

**Figure 1:**
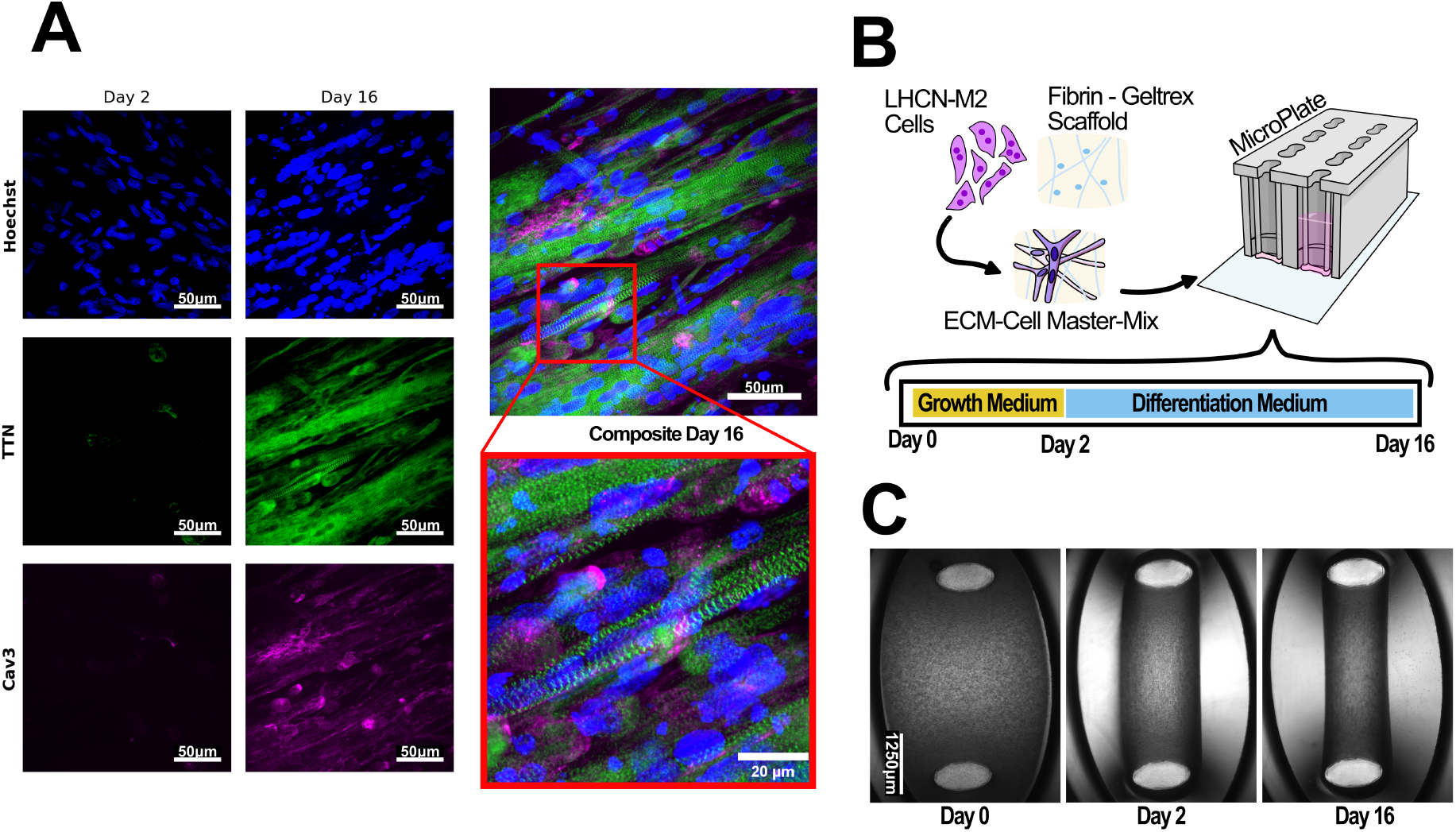
**A** maximum intensity Z-projections of a 24 *µm* thick confocal recording of LHCN-M2 engineered skeletal muscle tissues at Day 2 (after two days in growth medium) and at Day 16 (after additionally 14 days in serum-reduced differentiation medium). The tissues were immuno-stained for nuclei (Hoechst, blue), the sarcomeric protein titin (TTN, green), and the myogenic marker caveolin-3 (CAV3, magenta). At Day 16, the composite image and high-magnification inset demonstrate the transition from mononuclear myoblasts to multinucleated, aligned myofibers exhibiting highly ordered, periodic TTN striations. Tissues were imaged using a 60x water-immersion objective. **B** Schematic representation of the seeding procedure. LHCN-M2 cells are mixed with a fibrin-geltrex hydrogel and seeded in a specialized well-plate system facilitating two pillars per well. After 2 days in growth medium, cultures are switched to differentiation medium for 14 days to promote myotube formation. **C** Brightfield time-lapse imaging of a representative tissues. mages were captured immediately after seeding (Day 0), after 48 hours (Day 2), and after 16 days (Day 16). Rapid compaction occurs within the first 48 hours, resulting in a dense, cohesive tissue with preserved structural integrity over 16 days.

For each ESM 250.000 LHCN-M2 cells were mixed with a fibrinogen-geltrex scaffold. The polymerization of fibrinogen into fibrin was triggered by the addition of thrombin and immediately afterwards the suspension was casted into the wells of the MicroPlate. After 5 Minutes of incubation time at 37°C and 5%CO_2_ the lid was attached, submerging the pillars into the polymerized hydrogel to provide mechanical anchor points for the tissues. Each well was then filled up with 200 *µl* seeding medium.

Within the first 48h after casting the tissues, a prominent compaction of the hydrogel was observed as shown in Figure 1 C, indicating active cellular tension and force generation during the remodelling phase.

By day two the tissues also exhibited an anisotropic orientation along the longitudinal axis between the pillars. To induce the differentiation, the medium was switched to a growth-factor-reduced differentiation medium on the second day, which was then exchanged every other day for 14 days as illustrated in Figure 1 B.

To confirm the transition from proliferating myoblasts in to mature multinucleated myotubes, we performed immunofluorescent analysis on the second day (before switching to differentiation medium) and after additional 14 days in differentiation medium as shown in Figure 1 A. The tissues were fixed with a 4%PFA solution and then stained for Nuclei (Hoechst), Titin (Ttn) (sarcomeric alignment) and Caveolin-3 (CAV3) (sarcolemma maturation).

Confocal imaging showed that the myoblasts successfully fused into elongated multi-nucleated fibers. Titin expression displayed prominent striations, confirming the organization into sarcomeric units. Furthermore, the presence of the expression of CAV3 along the fibers indicates the development of the sarcolemma, which is important for a functional excitation-contraction coupling (see Figure 1 A).

### Microtissues are electrically excitable and show reproducable force generation

Beyond structural alignment, we assessed the functional physiological state of the ESMs by monitoring contractile activity. Given the stiffness of the PMMA pillars 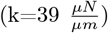 (6) contractile events were expected to produce deflections on the micrometer scale. These contractions are barely visible in 2X-brightfield images and subtile, but clearly visible with 10x maginifcation (see Figure 2 A). By using a custom high-resolution tracking pipeline (see Methods and (30)), these deflection can be translated into high-fidelity contraction curves to evaluate the functional performance of the tissues, thus providing the raw data necessary for subsequent kinetic modeling.

**Figure 2:**
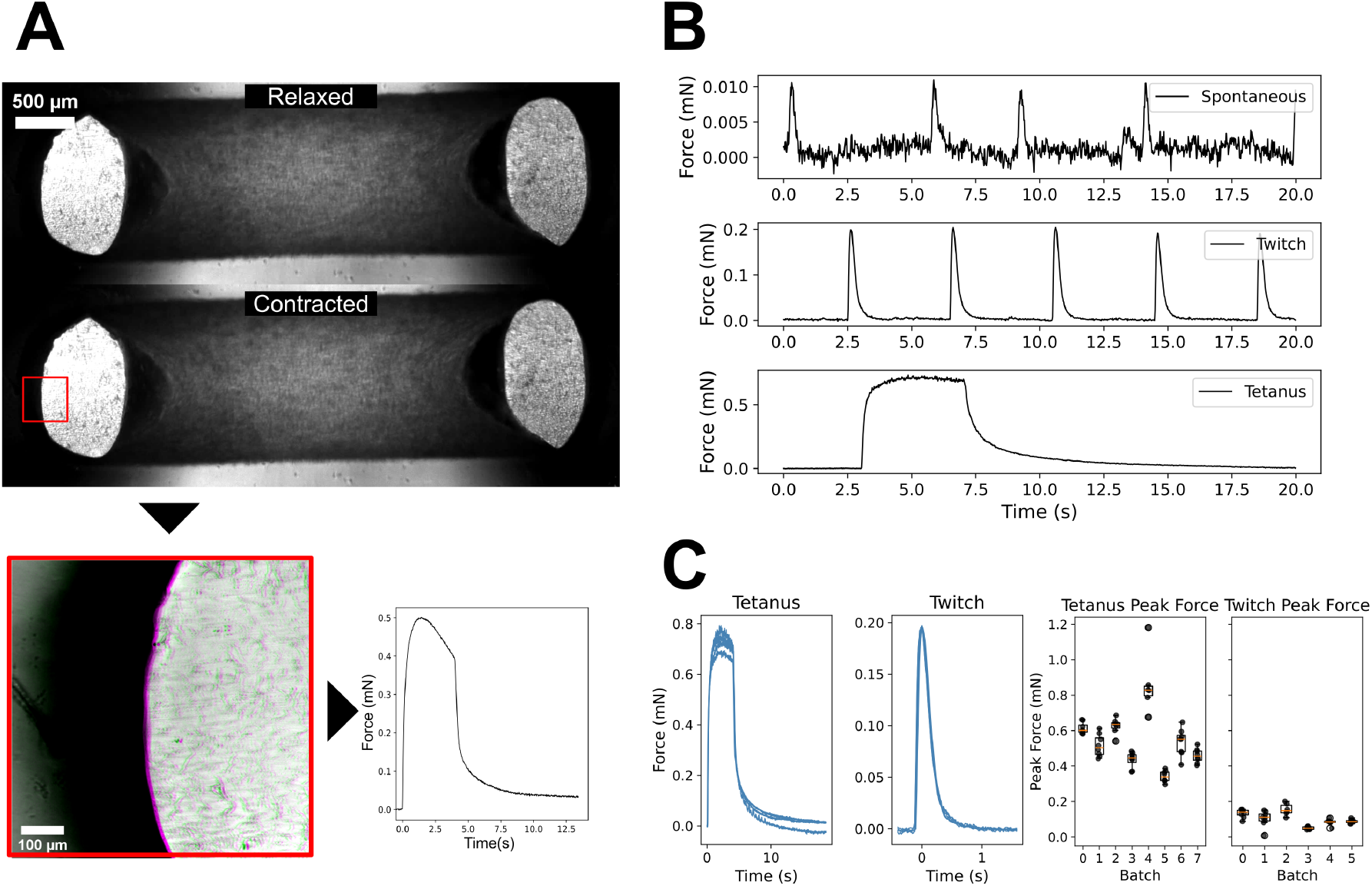
**A**: Representative images illustrating the methodology for force derivation via pillar displacement. While macroscopic 2x brightfield images show minimal perceptible movement between relaxed (upper) and contracted (lower) states, high-resolution 10x brightfield overlays (bottom) resolve the subtle deflection of the flexible posts (highlighted in magenta). This displacement is computationally translated into high-fidelity contraction curves (force vs. time) with a python-based analysis pipeline, enabling the quantification of contractile force. **B**: Representative contractile responses obtained from the same tissue: unstimulated spontaneous rhythmic contractions (top), short-duration twitch contractions (middle), and sustained tetanic contractions (bottom). **C** (left) Superimposed contraction-curves of repeated tetanus and twitch cycles within the same tissue highlight high functional stability and intra-sample reproducibility. (right) Boxplots quantifying peak force for different tissues across multiple independent batches. For tetanus peak forces a moderate batch-to-batch variation (*σ*_*inter*_ = 15%) is observed while the variation within a single batch is rather small (*σ*_*intra*_ = 7%). For twitches, both the inter-batch variation as well as the intra-batch variation is rather low (*σ*_*inter*_ = 3.6%, *σ*_*intra*_ = 2.6%).

Notably, several tissues exhibited detectable spontaneous contractions as shown in Figure 2 B, with magnitudes in the range of 0.1 − 0.4*mN* . These spontaneous events are potentially driven by rhythmic calcium oscillations, are considered as an indicator of advanced functional maturation and were also reported in other studies (12, 31, 32). Such activity suggests that the myotubes have developed a polarized sarcolemma and a mature sarcoplasmic reticulum capable of autonomous, cyclic calcium handling in the absence of external triggers (9) .

While spontaneous activity confirms biological viability, the resulting contractions are uncontrolled. To achieve the synchronized and comparable kinetics required for a rigorous quantitative analysis, we implemented an electrical stimulation protocol using steel electrodes positioned adjacent to the pillars within the culture well.

Brief pulses (5 V bipolar, 0.1 s duration at 100Hz switching between ±5V) successfully evoked discrete twitches. To assess the maximum contractile capacity, we applied high-frequency stimulation (±5V bipolar, 4 s duration at 100Hz) for 4 seconds. The resulting force-time profiles shown in 2 B exhibited a characteristic rapid onset followed by a stable, sustained plateau. This response confirms the presence of a functional excitation-contraction coupling machinery capable of maintaining force production under high-frequency depolarization.

Repeated stimulation of individual tissues yielded nearly identical force profiles for both twitch and tetanus events. Furthermore, an analysis of peak force revealed minimal inter-tissue and moderate inter-batch variation (Tetanus: *σ*_*inter*_ = 15%, *σ*_*intra*_ = 7%, Twitch: *σ*_*inter*_ = 3.6%, *σ*_*intra*_ = 2.6%) (see Figure 2 C), establishing the LHCN-M2 ESMs as a robust and reliable system for longitudinal pharmacological studies and biophysical modeling.

### Contractile properties of LHCN-M2 can be modified by addition of pharmacological substances

The generation of contractile force is the functional hallmark of skeletal muscle, emerging from a highly coordinated biochemical cascade. To validate the LHCN-M2 ESMs as a robust platform for biophysical force measurements, drug discovery and disease modeling, we interrogated three distinct functional tiers: the Troponin-Calcium machinery, the Actomyosin cross-bridge cycle, and the structural cytoskeleton. We perform the force generating analysis on the tetanus contractions. This ensures that we asses the maximum force generation capacities of the muscle, and to get a clear separation between the contractile phase, the force plateau and the relaxation phase.

To quantify the effect of the different pharmacological and cytoskeleton affecting compounds, each tissue was subjected to three tetanic contractions before compound application to obtain a baseline measurement (yellow in Figure 4 B). This was followed by a 120-minute incubation with either the pharmacological agent or a DMSO vehicle control, and again three tetanic contractions (blue in Figure 4 B). To take into account tissue- or batch specific variations we performed a post-treatment/pre-treatment normalization strategy. To ensure a stable pH environment during extended pharmacological assessment, the culture medium was supplemented with HEPES (10 mM). Statistical significance was determined using the Mann-Whitney U test, comparing the normalized force change of the drug groups against the DMSO control. Interestingly, the DMSO control group exhibited a consistent, albeit slight, increase in peak force over the 120-minute window (Figure 4 A), potentially reflecting structural realignment and improved fiber recruitment following the initial contractions.

To assess the platform’s sensitivity to calcium-sensitizing agents, we applied the fast skeletal muscle troponin activators (FSTAs) Tirasemtiv (33) and Reldesemtiv (34), that enhance calcium binding affinity to Troponin C (See also Figure 3). In accordance with previous findings in murine muscle models (34–36), both compounds showed a marked increase in force generation at sub-maximal activation levels (see SI Figure S1). At concentrations of 10 *µM*, both drugs significantly elevated peak force output in tetanus contractions, while intermediate concentrations (1 *µM*) produced graded responses (Figure 4 A), confirming dose-dependent potentiation of the calcium-sensitivity machinery and validating the platform’s suitability for contractility-enhancing drug assessment.

**Figure 3:**
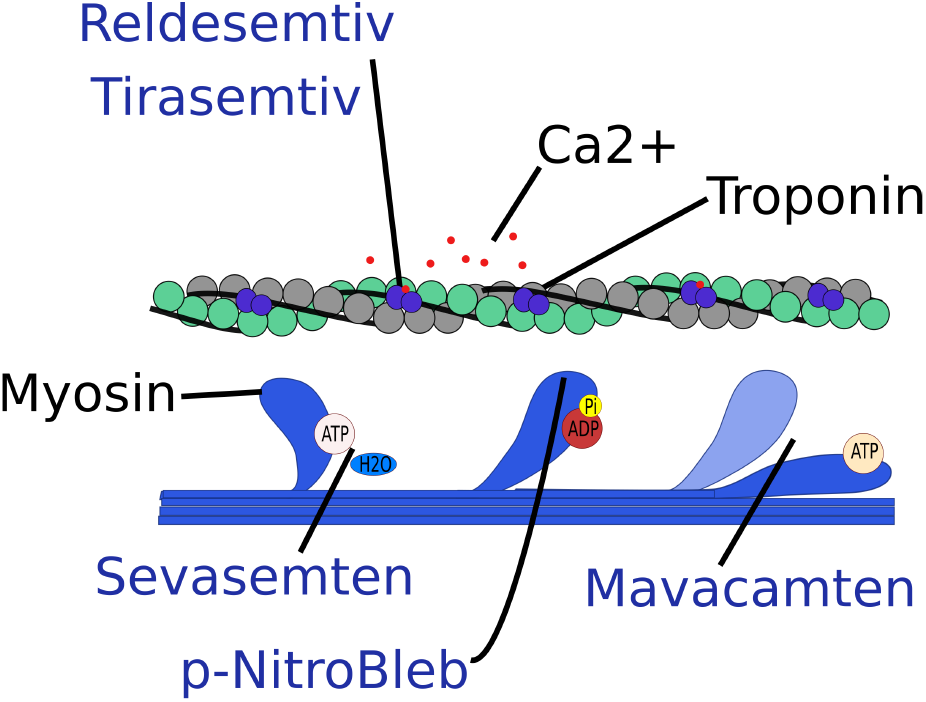
Schematic illustration of the drugs affecting the cross-bridging. Relsedemtiv and Tirasemtiv sensitize troponin to Ca2+. Sevasemten reduces the hydrolyzation of ATP. Para-nitro-blebbistatin prevents attachment of myosin heads to the actin. Mavacamtem transfers myosin heads into the super-relaxed state

To evaluate modulation of the cross-bridge cycling, we administered the small-molecule myosin inhibitors p-Nitroblebbistatin, Sevasemten, and Mavacamten. P-Nitroblebbistatin - a non-phototoxic, non-cytotoxic derivative of blebbistatin with similar inhibotory properties(37) - retains potent inhibition of class II myosins through binding near the nucleotide binding site at the myosin lever arm (38). Dose dependent effects of Blebbistatin on the contractile force were previously reported in cardiac tissues at concentrations of 2*µM* (39). Similar experiments in C2C12 microtissues showed complete abolition of contractility at concentrations of 100*µ*M (7). In our experiments we observed that 100*µ*M indeed had a strong effect on the peak force by reducing it to 40% of its initial peak force (Figure 4 A). However, the overall shape of the curves and i.e. the ability to sustain the plateau were rather unaffected (Figure 4 B). p-Nitroblebbistatin is expected to target non-muscle myosin so that could also affect the viscoelastic properties of the tissue.

**Figure 4:**
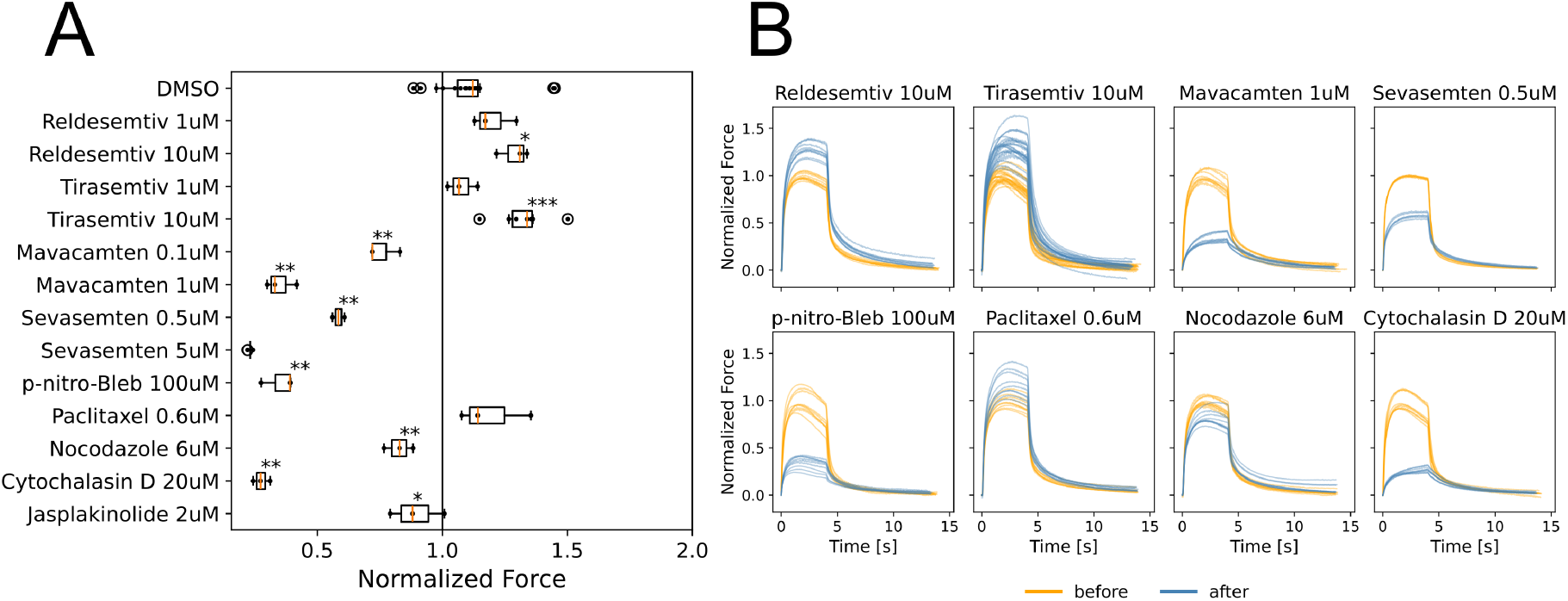
**A** Effect of nine pharmacological compounds on the peak contractile force of the engineered tissues alongside a DMSO vehicle control. To account for inherent biological and experimental variance, force measurements were recorded as the mean of three consecutive tetanic contractions per tissue both prior to and after a 120-minute incubation period. Data are presented as normalized peak force relative to the pre-drug mean of each respective tissue. Statistical significance was determined using the non-parametric Mann-Whitney U test, comparing the effect of each treatment group (*N* ≥ 3 except for Sevasemten 0.5*µM* N=2) against the DMSO vehicle control (*N* = 19). The results demonstrate the platform’s sensitivity in detecting both potentiation and inhibition of contractile force. **B** Representative contraction curves illustrating functional changes of different pharmacological compounds beyond peak force modulation. For each drug, the pre-drug contraction (orange, normalized to the mean) and post-drug contraction (blue, normalized to the same pre-drug mean) are overlaid.

Mavacamten, originally developed as a selective cardiac myosin inhibitor for hypertrophic cardiomyopathy, also engages skeletal myosin isoforms (40, 41), promoting a super-relaxed conformation through a distinct allosteric mechanism compared to p-Nitroblebbistatin (38). Consistent with this cross-reactivity, Mavacamten strongly reduced the peak force of the tetanus contractions to 35% if applied at 1*µM*, but also significantly affected the force generation at concentrations as low as 0.1*µM* (Figure 4 A).

Sevasemten, a selective fast-skeletal myosin inhibitor, produced a pronounced, dose-dependent reduction in contractile force. Peak force were significantly reduced to 58% at 1*µM* and nearly abolished at 10*µM* (Figure 4 A). While Sevasemten’s high selectivity for fast myosin has been reported, the magnitude of inhibition observed here likely reflects the elevated concentrations used relative to its half maximal inhibitory concentration (*IC*_50_ *<* 1*µM*), consistent with reports of partial cardiac myosin inhibition at ≥ 10*µM* in skinned fiber assays (42).

To probe structural components of the contractile apparatus, we treated tissues with agents targeting microtubules and actin filaments: Nocodazole, Paclitaxel (Taxol), Cytochalasin D (CytoD), and Jasplakinolide

Depolymerization of microtubules by 6*µM* Nocodazole significantly decreased peak contractile force to 83%, while stabilization by 0.6*µM* Paclitaxel produced a non-significant but opposite trend to 120% (Figure 4 A), suggesting that cytoskeletal dynamics contribute subtly to mechanical coupling, sarcomeric registry and force transmission.

As expected, 20*µM* of the potent inhibitor of actin polymerization Cytochalasin D induced a profound effect on force generation by reducing it to 27% of its initial peak force (Figure 4 A). Notably, while post-deflections were significantly diminished, the tissues still exhibited visible cellular contractions in brightfield microscopy videos (see SI V1). These findings indicate that CytoD disrupts cortical actin and extracellular matrix coupling while sparing portions of the sarcomeric network, consistent with previous findings in C2C12 tissue constructs(7, 43). Jasplakinolide showed no significant effect at the concentrations tested, potentially due to the high stability of established filaments in mature ESMs.

Comparisons of the normalized tetanus contraction curves for different pharmacological substances show that changes in the peak force is indeed the most prominent effect. However, additional effects can be observed by visually inspecting the curves (Figure 4 B). The fast skeletal troponin activators Tirasemtiv increases the peak force (1.07 at 1*µM* and 1.33 at 10*µM*), but also reduces the fatiguing behavior. Mavacamten and Sevasemten do not only reduce the peak force but also heavily impact the time scales of the contraction and relaxation. The myosin-2 inhibitor p-Nitroblebbistatin also affects the peak force in a similar range, but does leave the timescales mostly unaffected. These observations suggest that the compounds have a broader impact on the overall dynamics, that are not captured by the peak force alone. To better quantify the dynamical changes, it would be important to determine parameters that describe the increase, fatigue and relaxation dynamics. For this we developed a phenomenological mathematical modeling and parameterization approach which is described in the following section.

### Minimal parameter model provides a mechanical fingerprint

To quantify the complex contraction and relaxation kinetics of ESMs, we sought a mathematical framework capable of capturing physiological variability with a minimal set of interpretable parameters. Historically, muscle modeling has been dominated by two paradigms: Huxley-type and Hill-type models. While Huxley-based models excel at linking molecular cross-bridge mechanics to macroscopic phenotypes, they require fitting complex partial differential equations to derive the parameters (44) and high-fidelity data on sarcomere length transitions and calcium signaling to actually identify them (45). These metrics that are notoriously difficult to resolve in thick, 3D fibrin-based tissues and are usually not available in force experiments of 3D-ESMs. Conversely, Hill-type models utilize arrangements of springs and dashpots, which typically yield solutions based on standard exponential functions (46). However, we found that mono-exponential functions lack the degrees of freedom required to accurately fit the diverse contractile profiles observed in our 3D human tissues. Rather than increasing model complexity through the addition of discrete exponentials, that introduce challenges regarding numerical stability, parameter identifiability, and regularization (47), we propose the use of stretched exponentials *F* (*t*) = *F*_0_ · *exp*(− (*t/τ*)^*β*^) to describe both the contraction and relaxation phases.

The force generation in skeletal muscle is a multi-scale interplay between active processes like the calcium release from the sarcoplasmic reticulum, actomyosin cross-bridge cycling and passive viscoelastic properties of the extracellular matrix (ECM) or the cytoskeleton. The rate-dependent nature of these phenomena (44) suggests a system governed by multiple exponential decays. However, our 3D tissues comprise thousands of individual fibers with intrinsic heterogeneity, including varying maturation states, multinucleation levels, and sarcomeric organization. Instead of attempting to isolate a specific number of discrete exponential components—a task that is mathematically ill-posed and numerically sensitive to noise— the stretched exponential implicitly incorporates this intrinsic variability by modeling a continuous distribution of relaxation times within a single, robust framework (48).

The stretched exponential is a clean way to model disordered systems with heterogenious relaxation like our 3D microtissues.

As it can be seen in Figure 5 B a single exponential is not sufficient to capture neither the shape of the contractions nor the decay. In context of parameter efficiency one can see that already for a simple double Exponential of the form 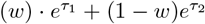 one would need one additional parameter for both of the contraction and relaxation phase compared to the streched exponentials 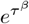.

**Figure 5:**
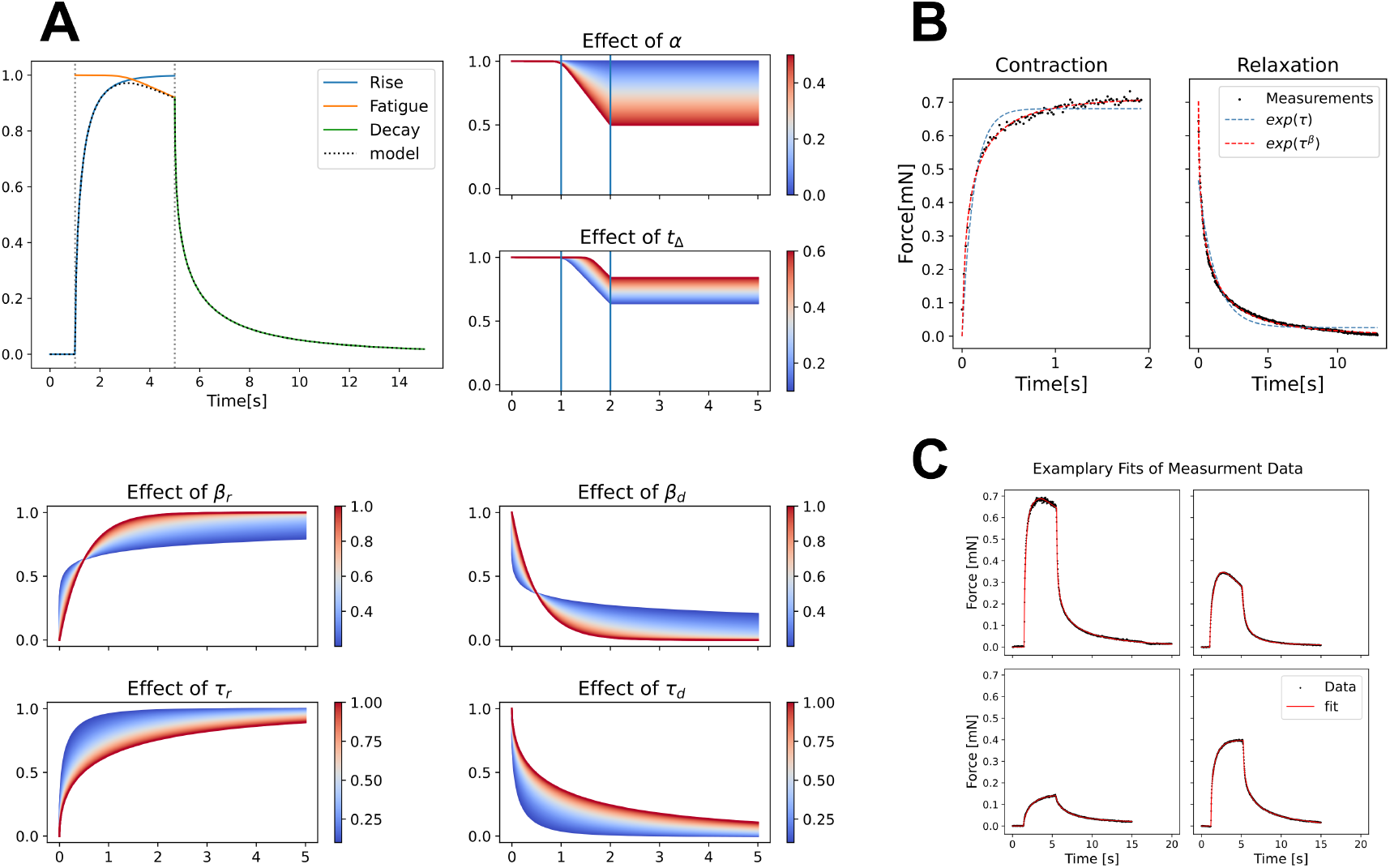
**A** Decomposition of the model into rise, fatigue, and decay components provides a 6 parameter model for the relevant dynamics. For each parameter its impact on the contraction curve is shown. *τ* modulates the global timescale while *β* dictates the “stretch” or asymmetry of the curve. Note the characteristic intersection point for varying *β* at a fixed *τ* . **B** Comparison between mono-exponential and stretched exponential to the rise and decay components of experimentally derived tetanus contractions. It shows superior accuracy of the stretched exponential over standard mono-exponential functions. Mono-exponential: RMSE_decay_ = 0.0196, RMSE_rise_ = 0.0296; Stretched-exponential: RMSE_decay_ = 0.0052, RMSE_rise_ = 0.0120 **C** Exemplary fits (red) of the stretched-exponential based model to drug-modulated contraction curves (black) demonstrate the good fit to a broad range of real measurement contraction curves.

**Figure 6:**
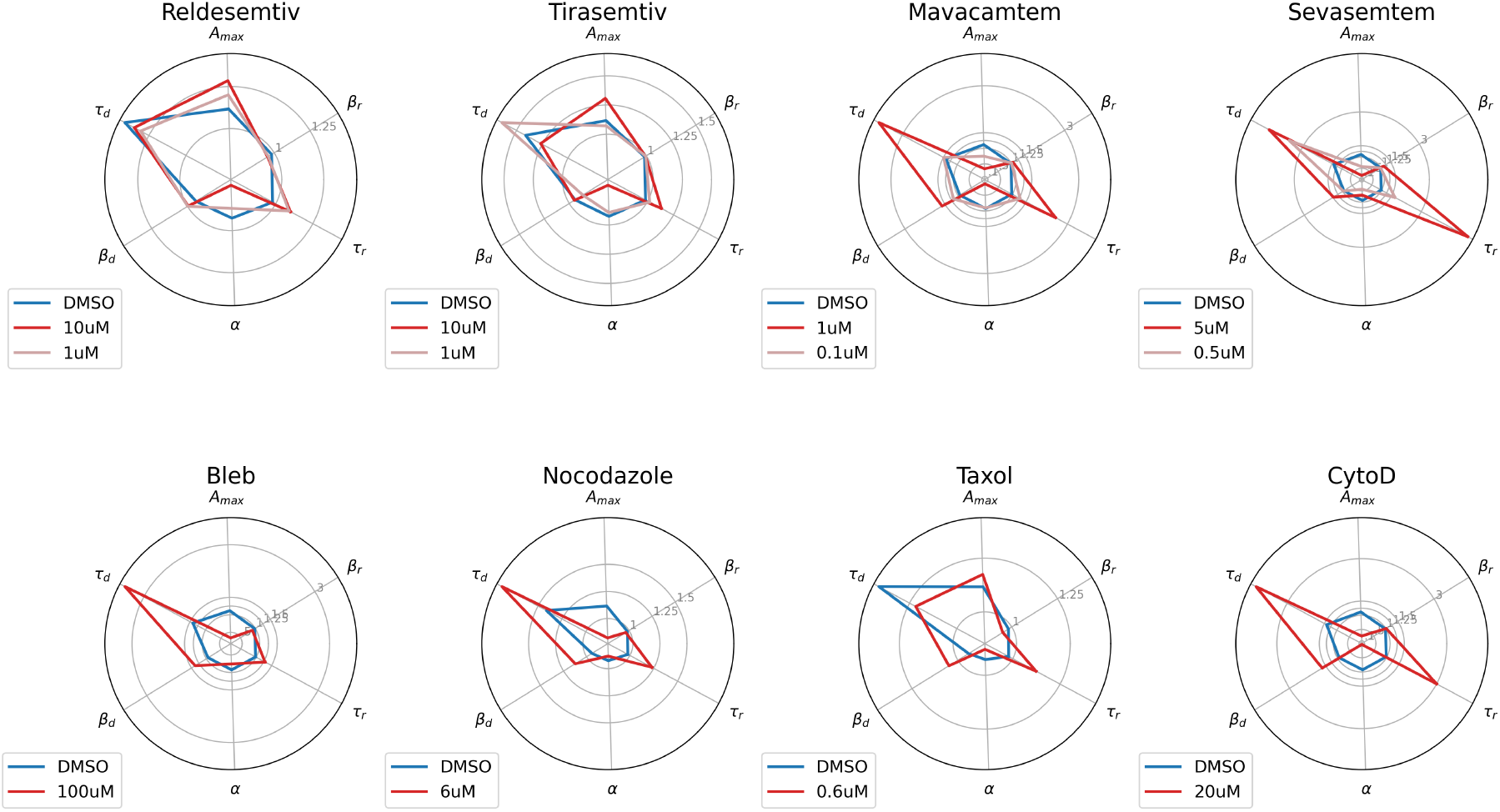
Effect of the different pharmacological compounds on the model parameters. All parameters have been normalized multiplicatively in respect to their value before the drug addition to compare the changes in value. DMSO control vehicle is plotted in blue for comparison. It can be seen that the drugs affect the contractions in different way and lead to distinctive profiles in the model parameters.

Instead, the proposed model consist of only 6 shape defining parameters in total, where two are used for each phase of rise, fatigue and decay. Most fundamentally the rise and decay of the contractions are each modeled with separate stretched exponentials. As seen in Figure 5 A the parameter *τ* serves as an effective timescale, conceptually analogous to the time constant in a standard mono-exponential model. It dictates the “half-life” or global rate of the process; specifically, it determines the point in time at which the curve reaches a fixed fraction of its asymptotic value. As *τ* increases, the entire contraction or relaxation profile is stretched along the time axis, representing a slower overall physiological response. Crucially, all curves sharing the same *τ* but differing in *β* intersect at a common characteristic point, the position of which is governed solely by the timescale.

The shape parameter *β* ∈ (0, 1] represents the “stretch” of the exponential and serves as a measure of the system’s kinetic heterogeneity. While *τ* sets the pace, *β* controls the asymmetry of the curve Figure (49). A value of *β* = 1 simplifies the model to a standard mono-exponential decay, representing a single-rate process. For *β <* 1, the model captures a process that is faster in its initial phase but exhibits a more pronounced “long-tail” behavior as it approaches the steady state 5 A. In the context of muscle engineering, this allows the model to capture e.g. the rapid initial cross-bridge recruitment followed by the slower, heterogeneous stabilization of the thousands of individual fibers within the 3D ECM.

The sustained phase of the tetanus is further refined by parameters governing the linear “fatigue” observed in several tissues. The slope parameter (*α*) and the onset delay (*t*_Δ_) allow the model to capture the gradual force loss during sustained high-frequency stimulation without interfering with the initial rise kinetics. The fatigue term acts multiplicative in the rise phase 5 A.

The absolute of the contractions is absorbed by a prefactor *A*. Additional parameters *t*_0_, *t*_1_ and *C* are incorporated to take into account the timepoints for the onset and offset of the electric stimulation and the Offset that can be caused by random drifts of the microscopy stage. These parameters are unrelated to the shape of the curve since they are merely shifting the curve according to our experimental setup. Thus they are not considered to be shape-defining parameters. Together this leads to the following model.

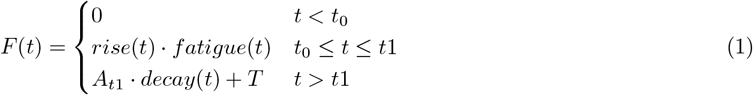

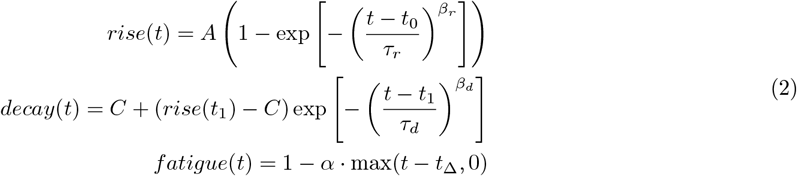

A description of the parameters can be found in Table 1

**Table 1:** Overview for the different model Parameters.

| Parameter | Function |
| --- | --- |
| $\tau_r, \tau_d$ | Time constants for rise and decay kinetics |
| $\beta_r, \beta_d$ | shape parameter (stretched exponential) |
| $A$ | Asymptotic maximum force amplitude |
| $t_0, t_1$ | Timepoints for start and end of the electric stimulation |
| $\alpha$ | Slope of the fatigue |
| $t_\Delta$ | Onset of the fatigue |
| $C$ | Offset |

To validate the ability of the model to generalize across a broad spectrum of contractile phenotypes, we applied the mathematical framework to the contraction curves derived from the pharmacological experiments. As demonstrated in Figure 5 C, the model yields excellent fits across a wide variety of contractile profiles, accurately capturing the diverse dynamics induced by the different compounds. Notably, even for heavily altered contractions such as those observed in Mavacamten treated tissues the model remains robust and maintains high fidelity. This capacity to characterize both baseline and pathologically or pharmacologically shifted kinetics indicates the strong generalization of the stretched exponential framework for human 3D muscle systems.

### Drug effect on model parameters

Thanks to the model, it is now possible to quantify the effect of the drugs in more detail by fitting the tetanus contractions across all drug-experiments. As explained above, we consider the 6 main parameters of *τ*_*r*_, *β*_*r*_, *τ*_*d*_, *β*_*d*_, *A* and *α* as the main relevant parameters, and can hence inspect the relative change for each parameter before and after the drug addition, as shown in Figure 6. DMSO is always plotted in blue for comparison, and was the same across all experiments as we had typically the same DMSO concentration in the applied drugs.

The most profound shifts in kinetic parameters (except for the near complete abolishment of high Sevasemten concentrations) were observed in tissues treated with Mavacamten. We found a robust, dose-dependent decrease in both the contraction rate (*τ*_*r*_) and the relaxation rate (*τ*_*d*_), while the shape parameters (*β*_*r*_*β*_*d*_) remained rather stable. This suggests that while Mavacamten significantly “passivates” the myosin heads, thereby slowing the transition into the force-generating state, the fundamental heterogeneity of the contraction process is preserved.

Similarly, Sevasemten significantly prolonged the relaxation phase. This pronounced effect on *τ*_*d*_ likely reflects a slowed cross-bridge detachment rate potentially mediated by e.g., altered cooperativity within the thick filament. In contrast, the fast-skeletal troponin activators, Reldesemtiv and Tirasemtiv, exhibited a distinct kinetic signature. At peak concentrations (10 *µ*M), both compounds slightly increased the decay speed. Since it was previously reported that FSTAs decrease the Ca-detachment rate (50) this suggest that the cross-bridge detachment or series-elastic recoil was the dominant rate-limiting step. Interestingly the two fast skeletal troponin activators are very similar profile in terms of model parameters, reflecting their shared mechanism of enhancing calcium sensitivity within the thin filament. Similarly, the two muscle myosin inhibitors Mavacamten and Sevasemtem produced consistent contractile patterns characterized by a distinct reduction in peak force and a marked alteration of the temporal parameters. While both drug classes modified the force output, their kinetic ‘fingerprints’ were qualitatively and quantitatively distinct from one another. This demonstrates that our model can distinguish between modulators of the thick (Actomyosin) and thin (Troponin) filaments

The model also revealed divergent effects for microtubule-targeting agents. Nocodazole and Paclitaxel exhibited opposing influences on the relaxation rate. While Nocodazole treatment led to slower relaxation kinetics, Paclitaxel appeared to accelerate the decay phase. This divergence suggests that microtubule stability directly influences the passive elastic recoil of the ESM; specifically, the stabilization of microtubules by Paclitaxel may increase the internal axial stiffness, thereby facilitating a faster return to the baseline state following tetanic contraction.

A key requirement for any analytical model in drug screening is the consistency of its parameters across experimental replicates. As shown in Figure 7, the parameter ranges for untreated tissues remained highly consistent across multiple independent batches. Notably, the *β*_*r*_ parameter behaved essentially as a constant across all conditions, suggesting it represents a fundamental structural characteristic of the LHCN-M2 myotube organization in the fibrin matrix. Even under pharmacological challenge, the *β* parameters stayed within a narrow, predictable range (Figure 7), while the time constants (*τ*_*r,d*_) exhibited high sensitivity. The observation that the strongest drug-induced variations occurred in the decay rate (*τ*_*decay*_) highlights the relaxation phase as a critical, yet often overlooked, indicator of muscle health and pharmacological response. By decoupling the “speed” of the contraction (*τ*) from its “shape” (*β*), our model provides a granular high-content readout that is highly reproducible across batches. Traditionally, contraction velocities are often calculated based on the time-to-peak or the one-half relaxation time (51–55).

**Figure 7:**
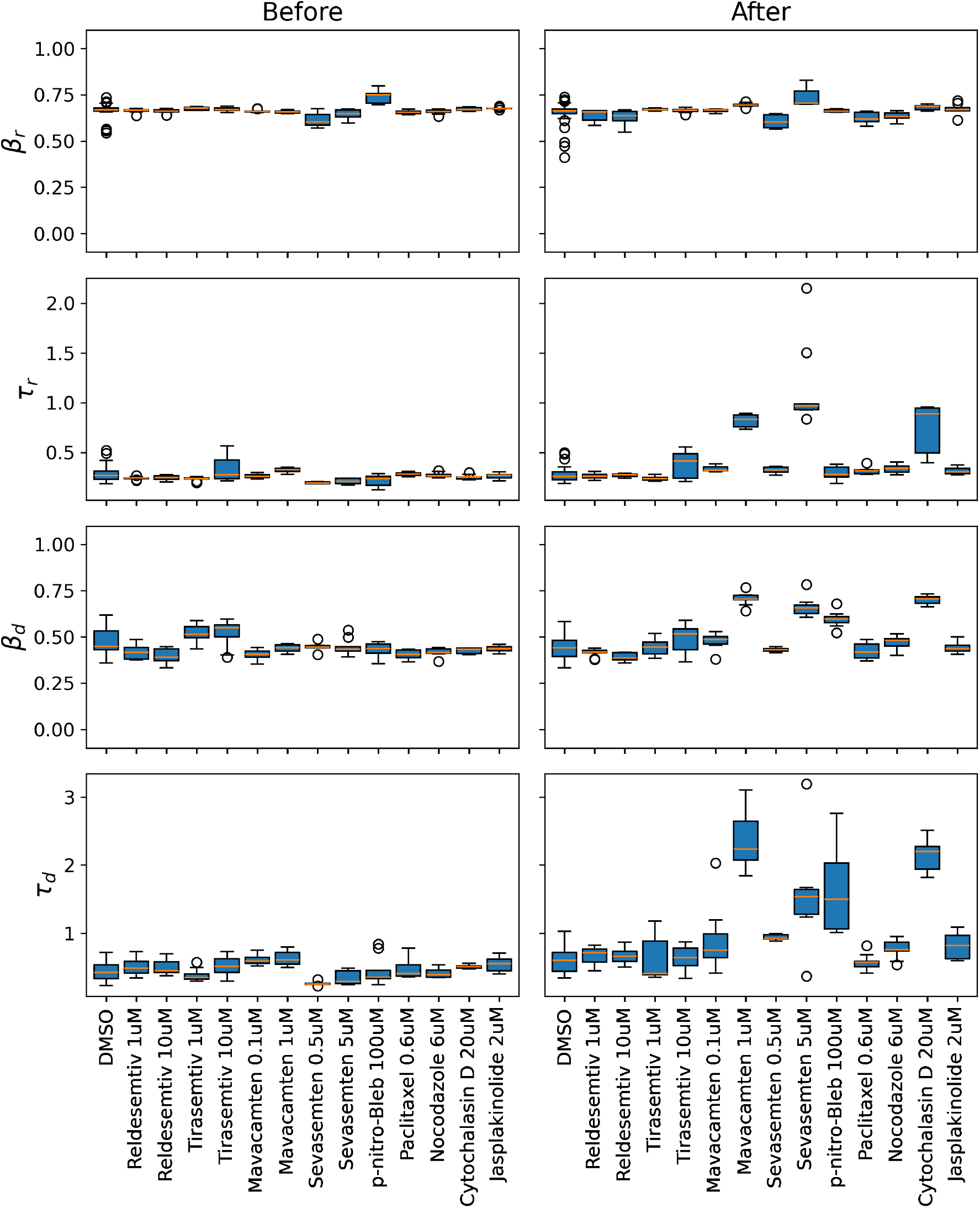
Boxplots for the shape defining parameters of the model (*β*_*r*_, *τ*_*r*_,*β*_*d*_ and *τ*_*d*_) before and after the addition of the pharmacological compounds. Before drug addition a high consistency across all batches and parameters and thus highlights the reproducibility of the LHCN-M2 microtissues. I.e. the *β*_*r*_ appears almost constant. After the drug addition the biggest effects are observed on the timescale of the decay *τ*_*d*_ especially for Mavacamtem, Sevasemtem and Cytochalasin D. These drugs also had the biggest impact on *τ*_*r*_ and *β*_*d*_, but on a smaller scale. *β*_*r*_ in contrast was only marginally altered even after drug addition.

Correlation analysis between heuristically derived velocities and the estimated timescales (*τ*_*r,d*_) of the stretched exponential confirmed that traditional metrics capture the global kinetic trends. However, the emergence of a clear non-linear relationship underscores that these heuristics oversimplify the underlying dynamics. In contrast the shape parameters (*β*_*r,d*_) seem to be uncorrelated with these traditional metrics. This demonstrates that the stretched exponential model captures additional layers of physiological data i.e. the ‘stretch’ or asymmetry of the curve, that are not contained within standard velocity measurements. By decoupling these parameters, our model offers a more granular and mathematically complete descriptor of muscle function.

### Proteomics Analysis reveals the expression of a broad set of differentiation markers and a hybrid fiber phenotypes

Finally, to support our pharmacological findings with molecular grounding, we performed proteomic analysis to validate the expression of specific drug targets and to characterize the developmental maturity of the engineered skeletal muscle tissues.

A global proteomic comparison between early-stage myoblasts (Day 2) and differentiated ESMs (Day 16) confirmed a successful transition from proliferating myoblasts to maturating myotubes. Hierarchical clustering (See SI Figure S5) revealed a distinct separation between conditions, reflecting a large-scale proteomic remodeling during the 14-day maturation period. Differential expression analysis, visualized via Volcano plot (Figure 8 A), confirmed a robust transition from a proliferative state to a committed myogenic lineage. This was marked by the significant downregulation of proliferation markers and a concomitant upregulation of the structural machinery essential for muscle function.

**Figure 8:**
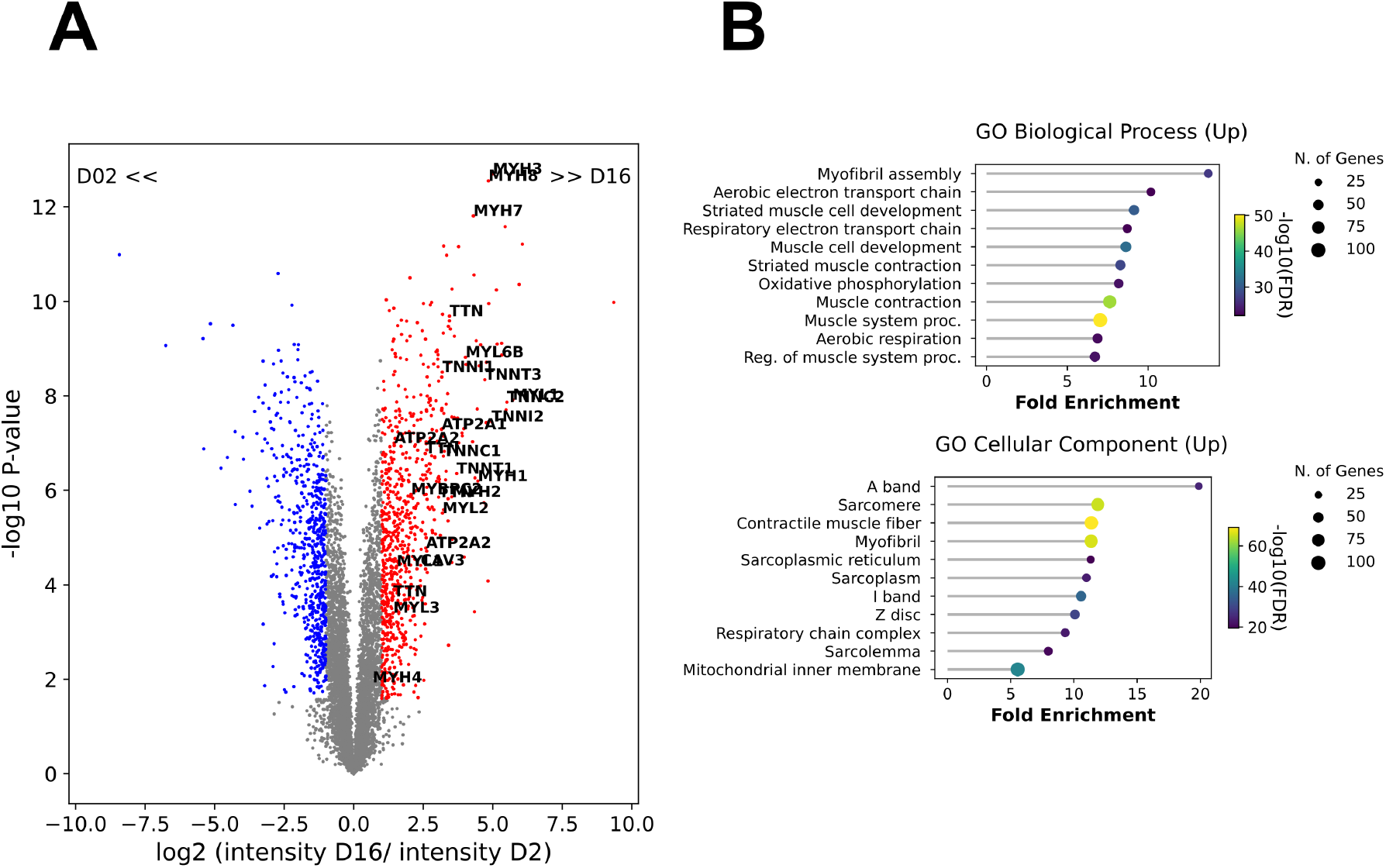
**A**) Volcano plot displaying the differential expression of the muscle proteome between Day 2 and Day 16 (*n* = 4 biological replicates per condition). The |log2| fold change (FC) is plotted against the (−log_10_) p-value for all identified protein groups. Statistical significance was defined using a False Discovery Rate (FDR) threshold of *q <* 0.05 and an effect size threshold of log2 FC *>* 1. Proteins are color-coded to indicate significant upregulation (red) or downregulation (blue) at Day 16, while non-significant proteins are shown in gray. Annotated hits highlight a significant enrichment of the contractile machinery and sarcomeric components, including multiple myosin heavy chain isoforms (MYH1, MYH2, MYH3, MYH4, MYH7, MYH8), titin (TTN), and the troponin complex (TNNC1/2, TNNI1/2, TNNT1/3). Furthermore, the upregulation of sarcolemmal CAV3 and calcium-handling ATPases (ATP2A1, ATP2A2) provides molecular validation for the functional maturation and Exciting-Contraction coupling observed in physiological assays. **B**): Lollipop plots illustrating the top enriched Biological Process and Cellular Component terms for proteins upregulated at Day 16. Terms are ranked by Fold Enrichment, with the bubble size indicating the absolute number of genes identified within each category and the color gradient representing the statistical significance (− log_10_ FDR). The Biological Process profile demonstrates a coordinated shift toward a mature contractile phenotype, highlighted by the significant enrichment of myofibril assembly and striated muscle contraction. A metabolic maturation is evidenced by the enrichment of the aerobic electron transport chain and oxidative phosphorylation pathways. The Cellular Component analysis provides a structural map of this maturation, showing high-magnitude enrichment for the sarcomere, specifically the A band, I band, and Z disc.

The differentiation process induced a significant enrichment of several Myosin Heavy Chain (MHC) isoforms, signaling the transition toward a mature contractile apparatus. While the presence of embryonic (MYH3) and neonatal (MYH8) isoforms reflects the expected in vitro developmental state (15, 55), we detected a clear emergence of the adult isoform for slow twitch/cardiac isoform (MYH7) as well as fast-twitch variants (MYH1, MYH2).

Gene Ontology (GO) enrichment analysis validated this maturation. We find strong upregulation concentrated in biological processes and components related to muscle development and contraction. Conversely, a downregulation of signatures related to cell-substrate adhesion, extracellular matrix remodeling, and cell migration confirms that by Day 16, the initial proliferative and migratory machinery has been silenced. As shown in Figure 8 B and Figure S4, the LHCN-M2 cells had successfully transitioned into post-mitotic, metabolically active, and pharmacologically responsive muscle fibers.

Next we correlated the proteomic abundance of specific targets with the observed functional shifts in model parameters to interpret the observed contractile phenotypes.

The robust expression of fast myosin heavy chains (MYH1 and MYH2) as well as myosin light chains, troponin and SERCA-pumps show the expression of fast-twitch related fibers (see Figure 8 A and Table S1) (56) and thereby the necessary molecular targets for Sevasemten (CK-3828136), which we already introduced as a selective fast skeletal muscle myosin inhibitor. However, the iBAQ quantification also shows a strong presence of slow-twitch related fibers (see Table S1) which seems insufficient to explain the near-total abolition of contractility observed at 10 *µ*M. This apparent discrepancy may be attributed to several factors inherent to engineered muscle tissues.

First, even adult *ex-vivo* fibers can occur in hybrid fiber-types without a clear fast or slow twitch myosin heavy chain profile (57) or re-express embryonic and prenatal isoforms for several weeks-during during regeneration(58). *In-vitro* myotubes are often still in the developmental state are thus expected to express the substantial shares of embryonic and prenatal myosin isoforms (MYH8, MYH3) that were detected by IBAQ analysis (see Table S1). It is possible that Sevasemten exhibits broader inhibitory activity across these immature or hybrid MHC variants. Second, previous studies in murine models have indicated that while Sevasemten is highly selective, it can exert off-target effects on cardiac myosins at higher concentrations (≥ 10 *µ*M) (42). Given that our tissues express significant levels of the “cardiac/slow” isoform MYH7, the 10 *µ*M dose, which is high relative to the compound’s *IC*_50_ of 0.2 *µ*M, may have induced a combined inhibition of both fast and slow-twitch machinery. Ultimately, this complex proteomic landscape, characterized by the coexistence of multiple myosin isoforms (MYH1, MYH2, MYH7, and prenatal variants MYH3, MYH8), directly supports our mathematical approach as we expect that different myosins can contribute different timescales. This intrinsic distribution of contractile proteins justifies hence the use of the stretched exponential *β* parameter, which is specifically designed to capture the resulting heterogeneity in contractile rates which a mono-exponential model would fail to resolve as it is exclusively modeling a single timescale. The strong upregulation of the fast-twitch related troponin complex (See Table S1), provides a direct target for the fast skeletal troponin activators Tirasemtiv and Reldesemtiv. The graded increase in force observed with these compounds is consistent with the high expression levels of their molecular substrate in the mature engineered skeletal microtissues.

## DISCUSSION

The development of the LHCN-M2 ESM platform represents a significant step toward scalable, human-relevant skeletal muscle modeling. By integrating 3D tissue engineering with a novel phenomenological modeling framework and proteomic profiling, we have established a pipeline that moves beyond the one dimensional peak-force analysis common in the field. Our findings demonstrate that the temporal “shape” of a muscle contraction represents a high-content readout that reveals the specific molecular compartment targeted by pharmacological agents.

### LHCN-M2 as a Scalable Alternative for Human Muscle Modeling

A major challenge in the field of muscle engineering is the trade-off between the scalability of murine lines (C2C12) and the translational relevance of human iPSC-derived tissues. While iPSCs are generally considered more physiologically, their use is often hampered by long differentiation protocols (often *>* 30 days) (15, 17) and high costs. Our results demonstrate that the immortalized LHCN-M2 line, when cultured in a 3D fibrin-Geltrex matrix, achieves significant functional maturation within 16 days. The structural registry confirmed by Titin striations and Caveolin-3 expression, coupled with the detection of spontaneous contractile activity, suggests a functional level of E-C coupling maturity.

Critically, our proteomic data confirms a global metabolic pivot, where the cells silence proliferative machinery to prioritize the assembly of the respiratory chain and the contractile apparatus. This confirms that LHCN-M2 ESMs are not merely a 3D cell culture but a developmentally committed tissue analog suitable for high-throughput screening.

### The Biophysical Significance of the Stretched Exponential Model

The introduction of a stretched exponential model addresses a critical analytical gap. While Huxley-type models provide deep biophysical insight, their large parameterization make it complex to fit the data in standard force-tracking data i.e. when sarcomere lengths and calcium transients can not be simultaneously resolved. Our model, requiring only six shape-defining parameters, offers a pragmatic yet powerful alternative. The mathematical choice of the stretched exponential is biologically motivated. The “stretch” parameter (*β*) effectively captures the heterogeneity of the tissue, thus reflecting a continuous distribution of relaxation times resulting from varying fiber maturation states and myosin isoform distribution. Our observation that *β*_*r*_ remains nearly constant across batches and drug treatments suggests it is a structural constant of the tissue’s architectural organization. In contrast, the high sensitivity of the time constants (*τ*_*r,d*_) to drug treatment validates them as the primary indicators of kinetic health. This decoupling of “heterogeneity” (*β*_*r,d*_) from “rate” (*τ*_*r,d*_) provides a level of granularity that standard Hill-type or mono-exponential models cannot achieve. In current practice, contraction and relaxation velocities are often estimated using non-parametric heuristics based on time-to-peak or the half-relaxation time. (51–55) These metrics implicitly assume a linear or fixed-rate process that cause a power-law relationship between these traditional markers and the *τ* parameters of the stretched exponential model. While these heuristics may capture general kinetic tendencies, their power-law, rather than linear, correlation confirms that they fail to accurately represent the underlying physiological rates across a broad spectrum of phenotypes.

Furthermore, these traditional metrics lack the robustness required for automated, large-scale analysis of complex phenotypes. For instance, in “fatiguing” muscle profiles, the half-relaxation interval frequently fails to isolate the relaxation phase from the preceding force decay, necessitating manual intervention or a-priori knowledge of the stimulation offset. By contrast, the joint model proposed here fits the entire contraction-relaxation cycle in a single, automated step. This eliminates the need for manual timepoint selection and provides a mathematically rigorous framework for characterizing complex, high-throughput contractile data.

### Decoding Pharmacological Mechanisms through Kinetic Fingerprinting

Our pharmacological results underscore that peak force is a relevant, but also coarse and potentially oversimplified metric for assessing muscle health and drug efficacy. While absolute force provides a snapshot of total contractile capacity, it fails to resolve the distinct molecular pathways through which different compounds operate. This is most evident when comparing the effects of Mavacamten, Sevasemtem, and para-nitro-blebbistatin, which are three compounds that all reduce peak force but induce different contractile patterns. Mavacamten’s profound slowing of the contraction rate (*τ*_*r*_), even at sub-micromolar concentrations (0.1*µ*M), supports its proposed mechanism of promoting a “super-relaxed” state. By effectively slowing their recruitment into the force-generating power-stroke, Mavacamten alters the entire temporal profile of the rise phase. In contrast, while p-Nitroblebbistatin also reduces peak force, it leaves the primary kinetic timescales relatively unaffected. This divergence highlights that myosin-targeting agents can have different “kinetic fingerprints”. Such differences would be entirely lost if the analysis relied solely on peak force

The divergent effects of Nocodazole and Paclitaxel on relaxation kinetics (*τ*_*d*_) provide new insights into the role of the non-sarcomeric cytoskeleton in muscle mechanics. The acceleration of decay by Paclitaxel suggests that stabilized microtubules act as internal “mechanical springs” that facilitate elastic recoil. This highlights the ESM as a tool not just for studying the motor proteins themselves, but for investigating the “mechanical circuit” of the whole fiber, including the ECM-cytoskeleton coupling that is often disrupted in muscular dystrophies (59).

### Proteomic Grounding of Functional Findings

The proteomic analysis provides the molecular justification for our functional observations. The significantly increased abundance of fast-twitch related proteins of the myosin heavy chain, myosin light chain, the sercapump and the troponin-complex explains the high sensitivity to Tirasemtiv, Reldesemtiv and Sevasemten. Interestingly, the detection of MYH7 (slow-twitch/cardiac myosin) explains why Mavacamten, a drug primarily optimized for cardiac tissue, exerted such a potent effect on our skeletal ESMs. The overall differential shift suggests that the cells have moved away from a proliferating phenotype towards a force-generating muscle phenotype.

### Limitations and Future Outlook

While the LHCN-M2 ESM platform demonstrates high reproducibility and pharmacological sensitivity, several limitations remain that provide avenues for future development. First, our proteomic analysis revealed the persistent expression of neonatal and embryonic myosin isoforms (MYH3 and MYH8). This indicates that, consistent with the majority of current *in-vitro* muscle models, the tissues remain in a developmentally immature state compared to adult native muscle. Future work could focus on increasing the degree of maturation through prolonged culture periods, more complex mechanical stretching (60, 61) protocols, or the integration of electrical stimulation (62) during the differentiation phase to drive the transition toward adult-type myosin profiles.

Furthermore, while this study focused on pharmacological agents targeting the active components of the contractile apparatus like the troponin-calcium machinery and the actomyosin cross-bridge cycle, contraction kinetics are also fundamentally governed by the tissue’s passive viscoelastic properties. Future studies should investigate the role of diverse ECM compositions (e.g., varying fibrinogen concentrations or the addition of collagen isoforms) and the modulation of scaffold stiffness on the resulting kinetic “fingerprints.” Experimentally verifying how internal viscosity and elastic recoil contribute to the stretched exponential decay would allow for a deeper understanding of muscle pathologies characterized by fibrosis or altered cytoskeletal rigidity. By combining our parametric modeling approach with precisely modulated ECM environments, it will be possible to distinguish between primary molecular motor dysfunction and secondary structural impairments.

Finally, a compelling next step would be to relate proteomic signatures to contractile phenotypes at a more granular level through targeted genetic engineering. The use of AAV-mediated gene delivery or CRISPR-based knock-in/knock-out strategies would allow for the precise modulation of proteins for which no small-molecule inhibitors exist. A primary candidate for such studies is Titin, as a critical sarcomeric protein acting as a molecular spring, its specific contribution to the elastic recoil and relaxation kinetics could be isolated. However, Titin is also a large protein and recent work showed that not all its genetic variation is well understood regarding cause and effect (63). Nevertheless, such functional genomics approaches, integrated with our stretched exponential framework, would enable a more direct mapping of the proteome to the physiological shape of muscle contraction, further enhancing the platform’s utility for complex disease modeling and precision medicine.

## CONCLUSION

We have presented a robust, integrated framework for the engineering and multiparametric analysis of human skeletal muscle tissues. By combining 3D engineered skeletal muscle tissue based on the LHCN-M2 cell line with a novel stretched-exponential modeling approach. We have demonstrated that it is possible to achieve reproducible contractile properties suitable for high-fidelity parametrization, enabling scaleable pharmacological screening. Our results prove that a drug’s effect is not just a matter of “how much” force is lost, but “how” the contraction dynamic is altered. This kinetic fingerprinting, validated by proteomics, provides a powerful new tool for understanding muscle biophysics and accelerating the preclinical development of myotropic therapies.

## MATERIALS AND METHODS

### Immunostaining

To evaluate structural maturation and sarcomeric organization, engineered skeletal muscle tissues (ESMs) were fixed and stained at two time points: day 2 (proliferative stage in growth medium) and day 16 (mature stage in differentiation medium).

The tissues were first washed with Phosphate-Buffered Saline (PBS) and subsequently fixed with a 4% paraformaldehyde (PFA) solution for 15 minutes at room temperature. Following fixation, the constructs were washed three times in PBS to remove residual PFA. To prevent non-specific binding and ensure membrane permeabilization, tissues were incubated in a blocking/permeabilization buffer containing 10% goat serum and 0.2% Triton X-100 for 1 hour at room temperature.

Primary and conjugated antibody staining was performed overnight at 4°C in the blocking solution. The following antibodies were used:

Sarcolemma maturation: Mouse monoclonal Caveolin-3 Antibody (A-3) conjugated to Alexa Fluor® 647 (Santa Cruz Biotechnology, 1:200).

Sarcomeric alignment: Titin Polyclonal Antibody conjugated to CoraLite® Plus 488 (Thermo Fisher Scientific, 1:200).

Nuclear visualization: Hoechst 33342 (Thermo Fisher Scientific, 1:500) was utilized to visualize multinucleation. Following overnight incubation, the tissues were washed three times in PBS to remove unbound antibodies. High-resolution fluorescent images were acquired using a confocal spinning disk microscope equipped with 20X and 60X water immersion objectives. To visualize the three-dimensional organization of the tissue, Z-stacks were captured and subsequently processed into maximum intensity projections. For high-magnification analysis of sarcomeric striations, a subset of the z-plane images were utilized to ensure optimal clarity and structural resolution. To facilitate a semi-quantitative comparison between undifferentiated (Day 2) and differentiated (Day 16) tissues, all imaging parameters, including laser power, exposure time, and detector gain,were maintained at constant settings across all samples.

### Cell Culture and Expansion

Immortalized human LHCN-M2 myoblasts were obtained from Evercyte (Vienna, Austria) and maintained according to the manufacturer’s recommendations with slight modifications. Cells were cultured in MyoUp growth medium, where 10% X-FBS was utilized as a substitute for the standard 15% Fetal Bovine Serum; no significant alterations in cellular morphology or proliferation rates were observed following this substitution. Prior to seeding, T75 culture flasks were coated with 0.1% gelatin for a minimum of 4 hours at 37°C. Myoblasts were initially plated at a density of 1,200 cells/*cm*^2^ and maintained in 20 mL of MyoUp medium. 10 mL of the medium was replaced every other day. To maintain a proliferative myogenic population, cells were passaged upon reaching 40% confluency. For 3D tissue generation, flasks were allowed to reach 80% confluency to ensure sufficient cell numbers.

### Fabrication of Engineered Skeletal Muscle Tissues (ESMs)

The 3D microtissues were fabricated using a modified version of a previously described protocol (6). For each construct, 250,000 LHCN-M2 cells were utilized. One day prior to tissue seeding, the polystyrene MicroPlates were sterilized via plasma treatment for 2 minutes using a Harrick plasma oven. Alternatively, sterilization can be achieved by 30 minutes of UV exposure. To prevent cell attachment to the well walls, each well was treated with 150 *µ*L of sterile-filtered 5% Pluronic F-127 solution to create a non-adhesive surface. The plates were sealed and incubated at 4°C overnight. The 3D scaffold consisted of a fibrin-Geltrex hydrogel matrix. Myoblasts were resuspended in a mixture of fibrinogen, Geltrex, and DMEM. Polymerization was initiated by the addition of thrombin, and the cell-hydrogel suspension was immediately cast into the prepared MicroPlate wells. Following polymerization, the CNC-milled PMMA lid, containing the cantilever pillars, was attached to provide mechanical anchor points.

### Differentiation and Maintenance

Following fabrication, the ESMs were maintained in growth-phase medium (MyoUp) supplemented with 3% aminocaproic acid (ACA) and 1% Penicillin/Streptomycin (P/S) for the first 48 hours. On day 2, differentiation was induced by switching to a medium consisting of DMEM and Medium 199 in a 4:1 ratio, supplemented with 2% Horse Serum, 4% ACA, and 1% P/S. The differentiation medium was exchanged every 48 hours throughout the 14-day maturation period.

### Proteomics

#### Sample Preparation and Lysis

LHCN-M2 ESMs (*n* = 4 per condition) were harvested at two time points: undifferentiated myoblasts (Day 2 in growth medium) and mature tissues (Day 16, following 14 days of differentiation). Tissues were washed, transferred to low-protein-binding tubes, snap-frozen in liquid nitrogen, and stored at -80°C. Samples were lysed in 1% sodium deoxycholate (SDC) and 100 mM HEPES using the BeatBox (PreOmics) homogenizer.

#### Protein digestion and MS Acquisition

Proteins were digested using the SP3 protocol with ReSyn Amine Beads on a Kingfisher Duo Prime platform. Peptide samples were spiked with Biognosys iRT standards for retention time calibration. Mass spectrometric analysis was performed on a Bruker timsTOF Pro 2 using data-independent acquisition (diaPASEF). Peptides (400 ng equivalent) were separated over a 100-minute gradient using an in-house 20x2 variable window method. Each biological replicate was analyzed in technical duplicate.

#### Bioinformatics and Statistical Analysis

Raw data were processed in Spectronaut v20.4 using the Pulsar search engine against the UniProtKB human reference proteome (v4.2025) plus a 54-protein contaminant database (1% FDR). Quantification utilized up to 6 fragments per peptide and 10 peptides per protein, with dynamic mass recalibration and quartile normalization. Statistical analysis was performed in Python. Technical replicates were averaged prior to a two-sided independent t-test. P-values were corrected for multiple testing using false discovery control (Q-values). Proteins with a log_2_ fold-change (FC) *>* 1 and *Q <* 0.05 were considered significantly regulated. One replicate was excluded from the final dataset due to low protein group identification. Unsupervised hierarchical clustering was performed using the Ward linkage method on Z-score normalized data to identify distinct protein cohorts based on relative abundance changes. Differential expression was visualized via volcano plots, mapping the log_2_ fold-change against the −log_10_ P-value. GO Enrichment Analysis was performed with the significantly up- or down-regulated proteins with the ShinyGO-Tool (64) based on the Gene Ontology (GO) knowledgebase (65, 66)

### Force Measurements

Functional assessment of the ESMs was performed by quantifying contractile forces via cantilever pillar deflection. To induce synchronized contractions, two stainless steel micro-electrodes (insect pins) were positioned within the well, posterior to the PMMA pillars, to deliver bipolar electrical stimuli. Contractile activity was characterized using two distinct stimulation protocols. Tetanic contractions were evoked by stimulating the tissues at an amplitude of ±5V with a pulse length of 10 ms (100 Hz) for a total duration of 4 s. This achieve a stable force plateau for the evaluation of maximum contractile capacity. Discrete twitch events were induced using an amplitude of ±5V and a pulse length of 10 ms (100Hz) applied over a duration of 0.1 s to assess individual contraction-relaxation cycles. Visual data were acquired using a Nikon Eclipse Ti2 microscope equipped with a 10X objective. High-speed video sequences were captured at 50 frames per second (FPS) (exposure time 20ms) using a Hamamatsu Orca Flash 4 camera. To maximize spatial resolution for the tracking algorithm, the camera’s Region of Interest (ROI) was centered on the distal edge of one cantilever post, capturing the micrometer-scale displacements during each contractile event.

### Force Analysis

Contractile force curves were derived by tracking the displacement of the PMMA cantilever pillars using a custom sub-pixel tracking algorithm implemented in Python (30). To capture these shifts, contractions were imaged via brightfield microscopy (10x magnification, 50 FPS) with the field of view centered at the distal edge of one pillar. The tracking was initialized by selecting a Point of Interest (POI) at the pillar-edge. To achieve sub-pixel precision and robust performance under varying imaging conditions the algorithm utilizes a stochastic template-matching approach. A two-dimensional Gaussian-shaped point cloud is sampled around the POI to interpolate pixel values, creating a smooth, weighted mask. The displacement (d) in consecutive frames was estimated by optimizing the spatial shift of this point cloud using a normalized negative cross-correlation (NNCC) metric combined with a gradient-free Cross-Entropy Method (CEM) (67, 68)optimization strategy.

The resulting displacement data were converted into contractile force (F) according to Hooke’s Law *F* = *k* · *d*), using a previously determined spring constant (*k* ≈ 39 N/m). To account for intrinsic tissue-to-tissue variability, forces were normalized to the respective average peak force obtained during baseline measurements prior to pharmacological treatment.

## AUTHOR CONTRIBUTIONS

M.L. and T.B. conceived and designed research; M.L. performed experiments with support from B.S.; M.L. prepared figures; M.L. drafted the manuscript with support from B.S. and T.B.; M.L. designed the model and the computational framework with support from T.B; M.L. implemented code; M.L. analyzed data; C.L. performed the proteomic workflow, M.L. and C.L. performed the statistical and bioinformatics analysis for the proteomics data; M.L. interpreted results of experiments with support from T.B. and C.L.; M.L. B.S., C.L. and T.B. edited and revised manuscript; M.L., B.S.,C.L., T.B. approved final version of manuscript.

## ACKNOWLEDGMENTS

Proteome analysis was supported by the infrastructure of the University Medical Center Göttingen (UMG) Core Facility Proteomics. Mass spectrometry equipment used in this study was jointly funded by the Deutsche Forschungsgemeinschaft (DFG) and the State of Lower Saxony under project 442069358. TB was supported by the Deutsche Forschungsgemeinschaft (DFG) under project 456112451, 569108445. This work was supported by the European Union’s Horizon Europe research and innovation programme under grant agreements No. 101123131 and No. 101247478.

## DECLARATION OF GENERATIVE AI AND AI-ASSISTED TECHNOLOGIES IN THE MANUSCRIPT PREPARATION PROCESS

During the preparation of this work the authors used Gemini in order to improve the language, flow, and clarity of the manuscript through proofreading and text formulation. After using this tool/service, the authors reviewed and edited the content as needed and take full responsibility for the content of the published article.

## SUPPLEMENTARY MATERIAL

**Figure S1:**
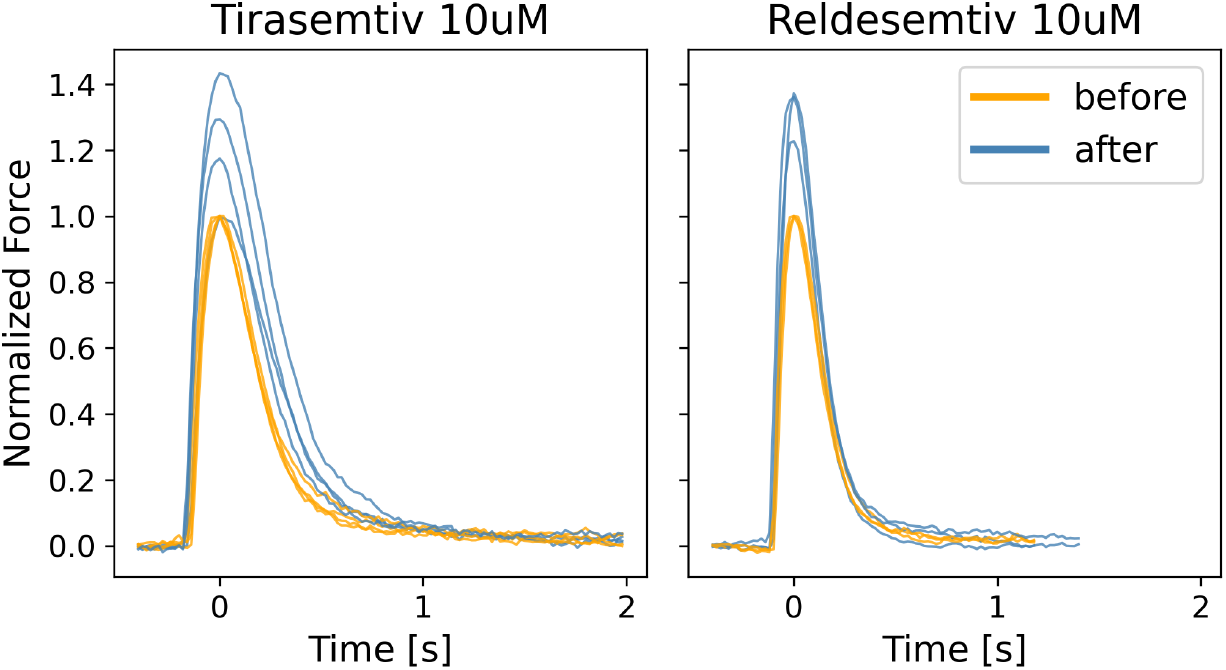
Normalized contraction curves for short twitches of tissues treated with Tirasemtiv (10*µ*M) and Reldesemtiv (10 *µ*M) show a force increase an submaximal stimulation

**Figure S2:**
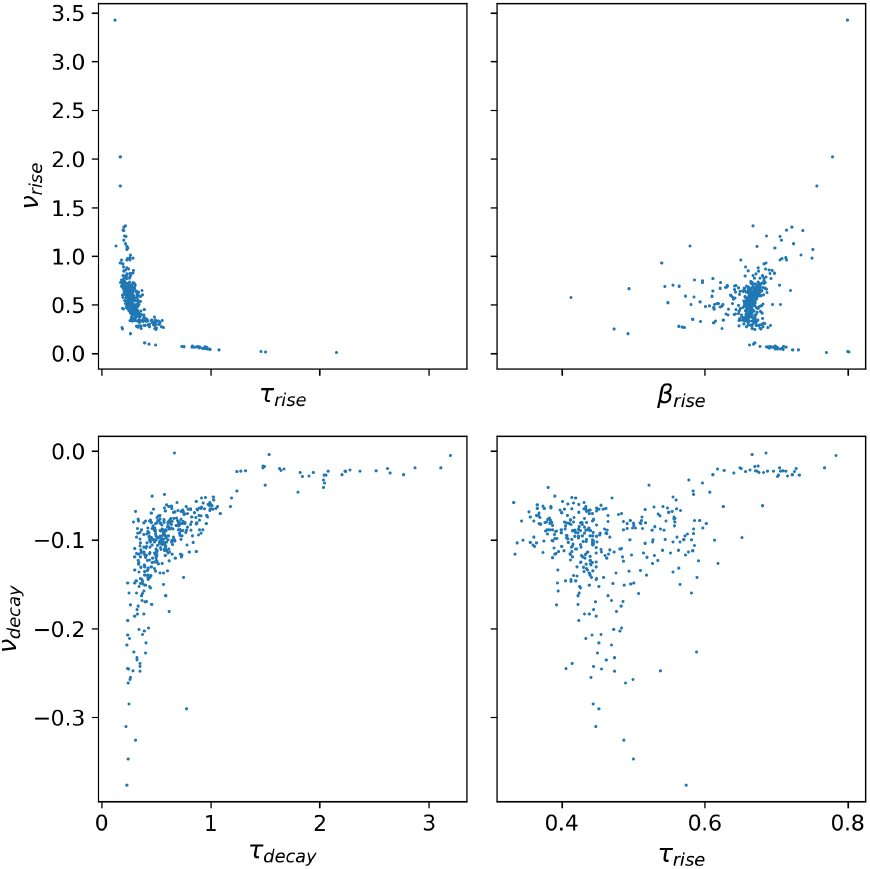
Correlation analysis between heuristically derived velocities **v**_*rise*_ **v**_*decay*_ and parameters of the stretched exponential *τ*_*r,d*_ and *β*_*r,d*_. While the beta parameters seem to be uncorrelated with the velocities, the tau parameters show a clear nonlinear relationship.

**Table S1:**
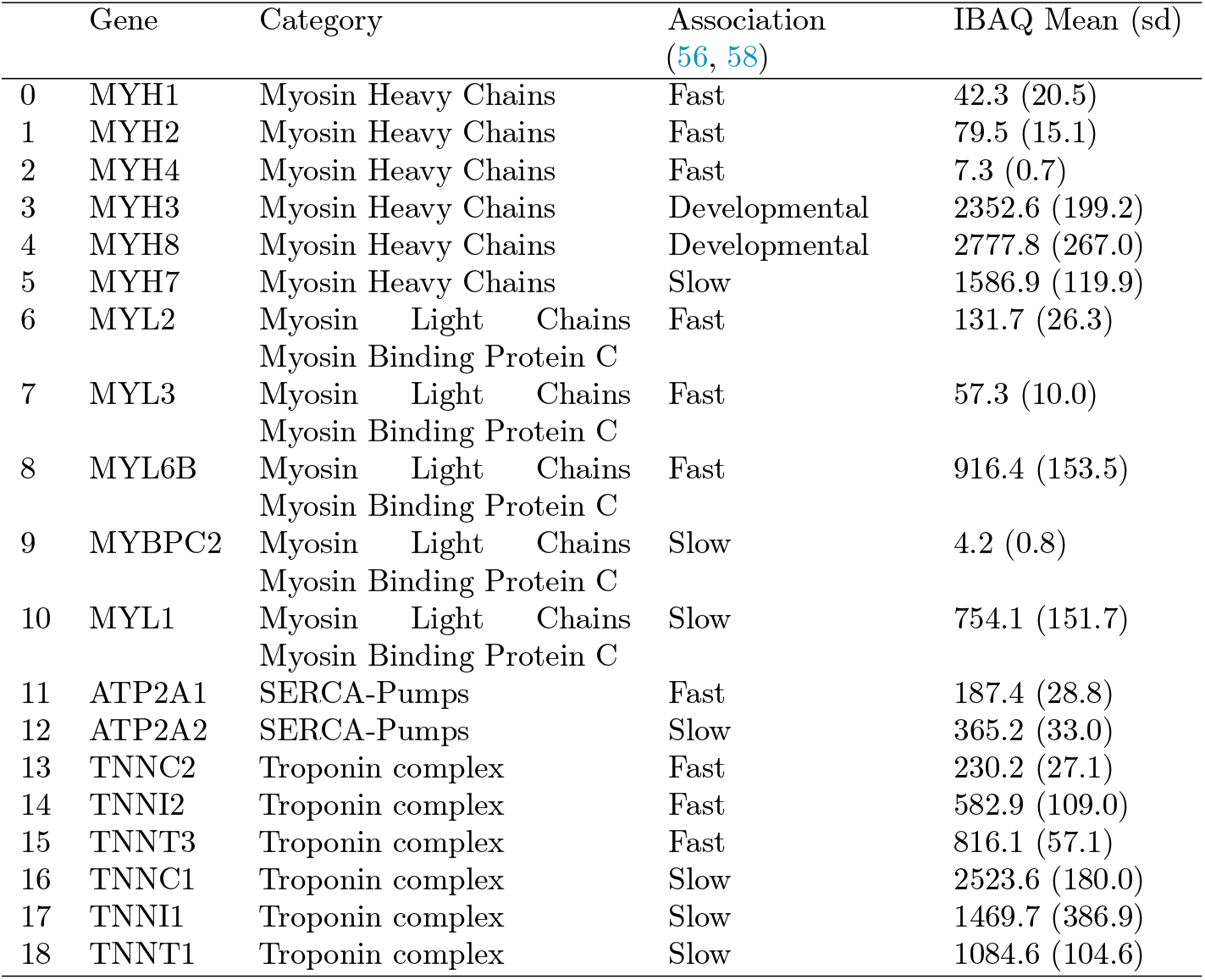
Overview of the proteomics data.

**Figure S3:**
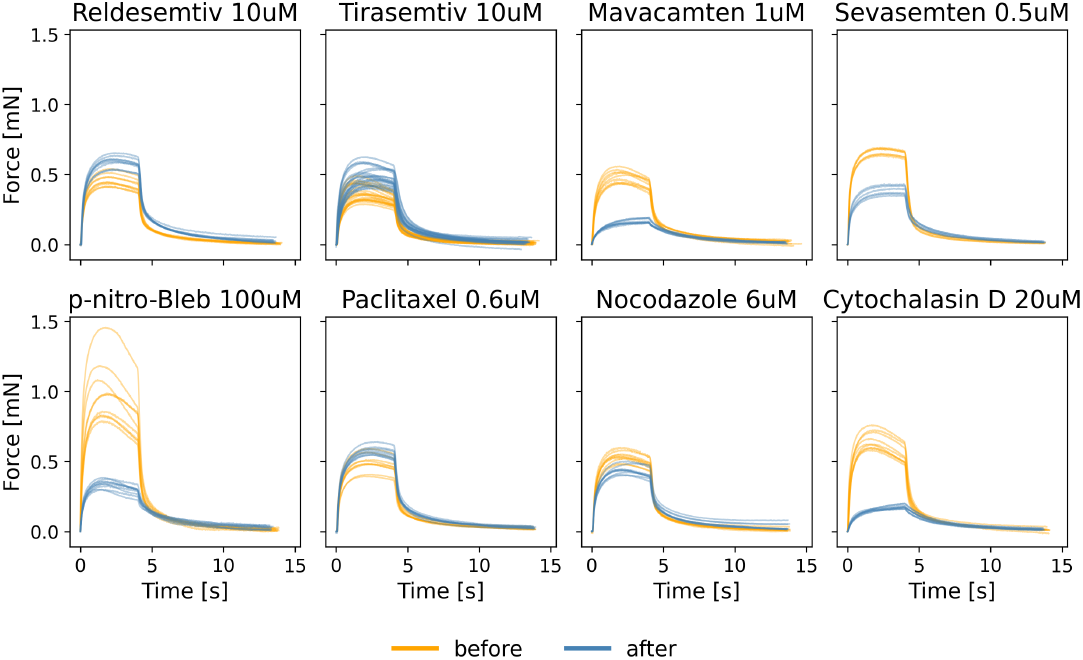
Impact of selected pharmacological compounds on the contraction curves.

**Figure S4:**
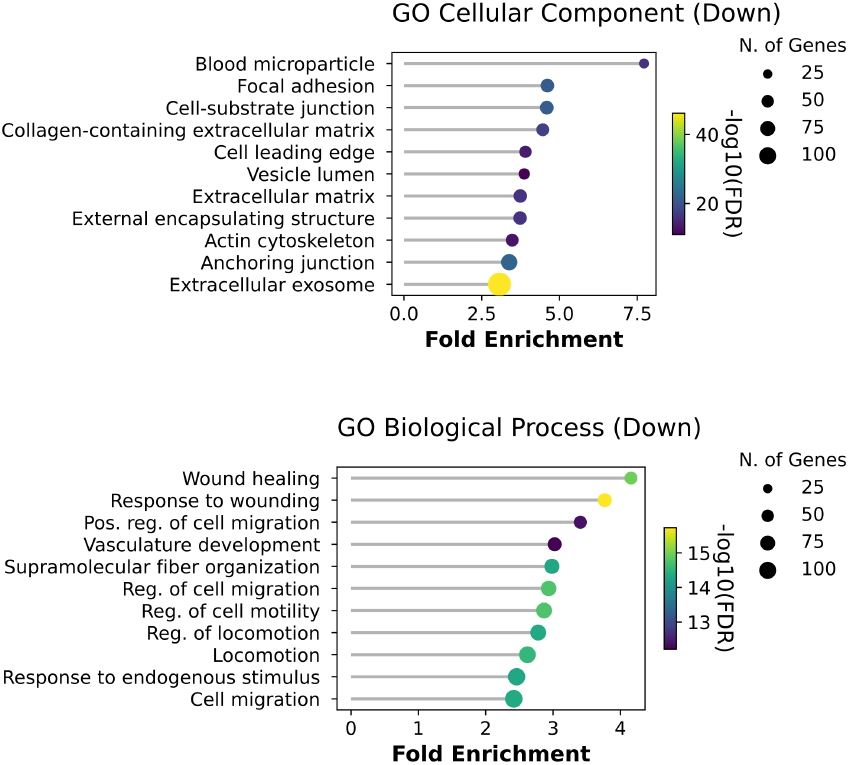
GO gene enrichment analysis for Cellular Components and Biological Processes that were significantly downregulated on day 16 compared to day 2. The Cellular Component analysis shows a significant decrease in proteins associated with the extracellular matrix, focal adhesions, and cell-substrate junctions. Correspondingly, the Biological Process analysis reveals the silencing of pathways related to cell migration, locomotion, and wound healing. Together, these data indicate a transition from a dynamic, migratory myoblast state to a stable, anchored myofiber phenotype.

**Figure S5:**
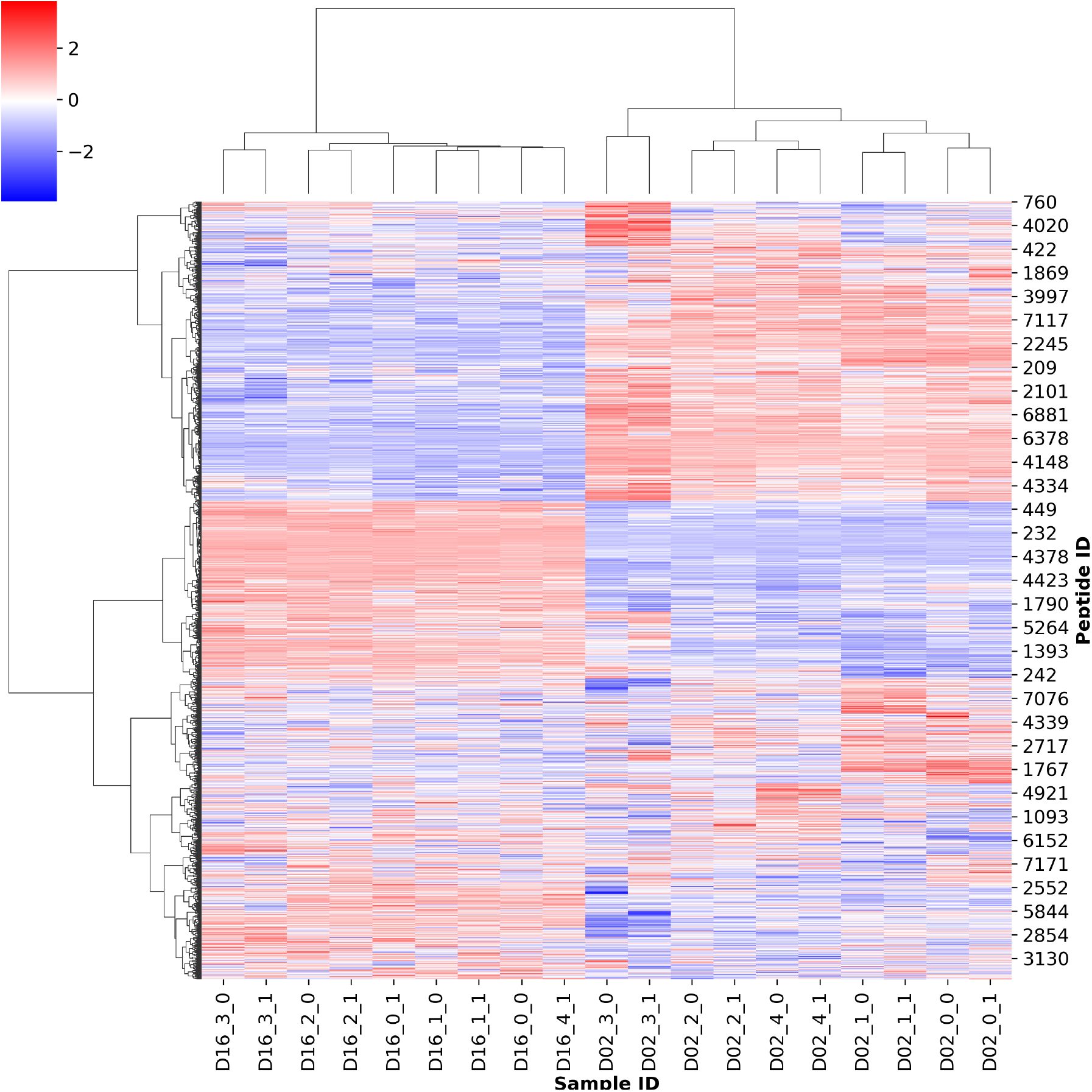
Unsupervised hierarchical clustering of differential protein expression. Heatmap and dendrogram illustrating the proteomic signature of hSMTs at Day 2 and Day 16. Data were normalized using a Z-score transformation across rows to emphasize relative abundance changes between time points. Clustering was performed using the Ward linkage method, minimizing within-cluster variance to identify distinct protein cohorts. The resulting clusters reveal a clear bifurcation of the proteome: a downregulated cluster (top) enriched in proliferative and migratory markers, and an upregulated cluster (bottom) dominated by contractile and metabolic proteins.

